# Laminar organization of encoding and memory reactivation in the parietal cortex

**DOI:** 10.1101/110684

**Authors:** Aaron A. Wilber, Ivan Skelin, Bruce L. McNaughton

**Affiliations:** Department of Psychology, Program in Neuroscience, Florida State University, Tallahassee, FL, 32308 USA; Canadian Centre for Behavioural Neuroscience, The University of Lethbridge, Lethbridge, AB, T1K 3M4 Canada; Department of Neurobiology and Behavior, University of California, Irvine, CA 92697, USA

**Author notes:** Contributed Equally (Aaron A. Wilber, Ivan Skelin).

**Keywords:** parietal cortex, posterior parietal cortex, modular organization, memory reactivation, multi-unit activity, high frequency local field potential, template matching

## Abstract

Egocentric neural coding has been observed in parietal cortex (PC), but its topographical and laminar organization is not well characterized. We used multi-site recording to look for evidence of local clustering and laminar consistency of linear and angular velocity encoding in multi-neuronal spiking activity (MUA) and in the high-frequency (300-900 Hz) component of the local field potential (HF-LFP), believed to reflect local spiking activity. Rats were trained to run many trials on a large circular platform, either to LED-cued goal locations or as a spatial sequence from memory. Tuning to specific self-motion states was observed consistently, and exhibited distinct cortical depth-invariant coding properties. These patterns of collective local and laminar activation during behavior were reactivated in compressed form during post-experience sleep, and temporally coupled to hippocampal sharp wave ripples. Thus, PC neuron motion encoding is consistent across cortical laminae, and this consistency is maintained during memory reactivation.

**Highlights:**

- Parietal cortex MUA encodes specific movements coherently across laminae.
- This organizational scheme is maintained during subsequent memory reactivation
- MUA and HF-LFP showed similar self-motion tuning and memory reactivation dynamics
- This establishes the utility of MUA and HF-LFP for human memory reactivation studies

## INTRODUCTION

A fundamental framework for neural coding in the parietal cortex is egocentric (e.g., Andersen et al., 1985; McNaughton et al., 1994; Nitz, 2006; Save et al., 2005; Save and Poucet, 2000; Schindler and Bartels, 2013; Whitlock et al., 2012; Wilber et al., 2014; Wolbers et al., 2008). In addition, some studies have also found evidence for allocentric (world-centered) encoding in parietal cortex (Chen et al., 1994a, b; Chen and Nakamura, 1998; Wilber et al., 2014). Subjective assessment of head direction tuning in parietal cortex suggested common tuning across depth on a given tetrode (Chen et al., 1994a, b; Wilber et al., 2014). The existence of larger organizational structure of these single cells (e.g., large scale populations of cells) has been postulated by the theoretical or computational approaches (e.g., Byrne et al., 2007; McNaughton et al., 1995; Xing and Andersen, 2000; Zipser and Andersen, 1988), but has not been empirically confirmed. Therefore, we set-out to look for evidence of population level organization of three types of previously reported ego-centric and allocentric reference frames, body centered cue direction (Wilber et al., 2014), self-motion tuning (McNaughton et al., 1994; Whitlock et al., 2012), and head direction tuning (Chen et al., 1994a, b; Wilber et al., 2014).

Behaviorally relevant neural activity patterns from single cells are reactivated during memory consolidation (Dupret et al., 2010; Lansink and Pennartz, 2014). Thus, we hypothesized that the functional relevance of modular organizational structure would be supported if the modular structure is also reflected in reactivation during post-experience-sleep – a postulated mechanism of memory consolidation originally demonstrated at the single cell level in hippocampus (Wilson and McNaughton, 1994) and subsequently shown for hippocampal-parietal interactions (Qin et al., 1997). There is now evidence that memory replay is tightly linked to plasticity at the level of single cells (Tavoni et al., 2015; van de Ven et al.; Yang et al., 2014). However, to our knowledge no study has looked for evidence of modular level reactivation.

Assessing the large scale population-level activity in animals would be useful for establishing the connection between animal microelectrode recordings and methods typically used for human neuroimaging electrocorticography (ECoG) and functional magnetic resonance imaging (fMRI), whose spatial and/or temporal resolution is usually limited to assessment of compound activity of large populations of cells. An increasing number of human ECoG studies provides a wide cortical coverage, but lack single neuron separability. Therefore, linking spiking activity with local field potential (LFP) features has been a growing area of research, most often finding the correlation between the spiking levels and LFP envelope in high-gamma range (Crone et al., 2011; Liu and Newsome, 2006; Ray and Maunsell, 2011); however, the high-gamma range includes synaptic current oscillations (Colgin et al., 2009; Fries, 2005) and not just spike-related currents. Therefore, we aimed to test whether the modular organization reflected in MUA is also detectable in high frequency LFP (>300 Hz), which to our knowledge has not been attributed any functional or physiological correlates other than spiking. The HF-LFP signal provides better temporal stability, relative to single neuron recording due to minor electrode drift, and therefore produces a more reliable readout for long-term studies and for brain machine interfaces (e.g., Gilja et al., 2012; Mazzoni et al., 2012). Similarly, recently an apparent connection between single unit recording studies in rodents (grid cells in rats; Hafting et al., 2005) to fMRI data in humans (Constantinescu et al., 2016; Doeller et al., 2010; Kriegeskorte and Storrs, 2016; Kunz et al., 2015), suggest that collective recordings may reveal a larger organizational structure of encoding that had largely been described at the single cell level in rodents (i.e., 6-fold symmetry initially observed in human fMRI studies during virtual navigation tasks). Thus, study of modularity of neural coding in animals using collective measures of local neural activity can help to bridge the gap between human and animal studies of neural coding and dynamics.

## METHODS

Male Fisher-Brown Norway hybrid rats (n=6), 5-10 months of age, underwent surgery for implantation of bilateral stimulating electrodes aimed at the medial forebrain bundle (MFB; 2.8 mm posterior from bregma, 1.7 mm from midline, 7.8 mm ventral from dura). Prior to surgery rats were housed 2-3 per cage. After recovery from surgery, rats were trained to nose poke for MFB stimulation. Then brain stimulation parameters (200 μs half cycle, 150 Hz biphasic 70-110 μA current applied for 300-450 ms) were adjusted to find the minimal intensity and duration that was sufficient for maintaining maximal responding. Next, rats with optimal MFB stimulation (N=3 of the 6 with stimulating electrodes) underwent surgery to implant a custom 18-tetrode bilateral “hyperdrive” or 18-tetrode unilateral hyperdrive aimed at the left parietal cortex (n=3; similar to: Kloosterman et al., 2009; Nguyen et al., 2009).

### Controls for MFB stimulation effects

MFB stimulation was necessary to obtain sufficient trials for some analyses. To ameliorate concerns about MFB effects on parietal cortex neural activity, data were removed for the brain stimulation duration plus an additional post-stimulation 20 ms blackout period (n=1; identical to: Kloosterman et al., 2009; Nguyen et al., 2009). The minimum post-stimulation blackout period of 20 ms was only applied for self-motion analyses, for memory replay analyses a longer (500 ms) blackout period applied because this was the time between trials when the cue light remained off for all trial and task types. In addition, MFB stimulation was delivered in one hemisphere and recordings were obtained from both hemispheres from most rats (n=2 of 3). For these rats we compared the proportion of tetrodes (pooled across sessions) that met the criteria for self-motion tuning in the same versus opposite hemisphere to brain stimulation. Similar to our previous report of no effect on proportion of any single cell types we measured (Bower et al., 2005; Euston and McNaughton, 2006; Euston et al., 2007; Johnson et al., 2010; Wilber et al., 2014), there were no differences in proportion of significantly self-motion tuned sessions (p<0.01) between hemispheres when self-motion tuning was measured with MUA (χ2_(1)_=2.29, p=0.13) and HF LFP signal (χ2_(1)_=0.28, p=0.60). Further, in previous experiments where it was possible to obtain sufficient coverage for analyses using food reward, identical results were obtained using MFB stimulation and food reward, suggesting that the results obtained from MFB experiments are generalizable (Euston and McNaughton, 2006). This suggests MFB stimulation did not directly influence the activity patterns we observed.

### Surgical Procedure

Recording arrays consisted of 18 tetrodes and 3-4 reference electrodes. Each tetrode consisted of four polyimide-coated, nichrome wires (12.5μm diameter) twisted together (Kanthal Palm Coast). The recording arrays were positioned over the parietal cortex (centered 4.5mm posterior from bregma and +/-2.95mm from midline) arranged in 1 or 2 closely packed guide-tube bundles with center-to-center tetrode spacing of ~300 μm. The arrays were positioned to target the average parietal cortex region for which we recently thoroughly characterized the connection densities (Mesina et al., 2016; Wilber et al., 2015) and also simultaneously record from the CA1 field of the hippocampus. These recording coordinates are likely to correspond to the intersection of the mouse anteriormedial, posteriormedial, and medio-medial areas (Wang and Burkhalter, 2007; Wang et al., 2012)

### Behavior

Training and testing took place on a large (1.5 m diameter) circular platform with 32 light cues evenly distributed around the perimeter (similar to the 8-station task; Bower et al., 2005). A custom computer program (interfaced with the maze via a field-programmable gate arrays (FPGA card, National Instruments) controlled maze events, delivered MFB stimulation rewards via a Stimulus Isolator (World Precision Instruments A365) when the rat entered a 10 cm diameter zone in front of the active cue light, and generated a coded timestamp in the Neuralynx system for each maze event (e.g., light onset for a particular zone). First, *alternation training* was achieved using barriers to restrict the movement of the rat to alternating between a pair of cue lights on opposite sides of the maze. To ensure cues were visually salient, lights were flashed at ~3 Hz (with equal on/off time) when activated. The first light was activated until the rat reached that reward zone and received MFB stimulation and then the cue light in the opposite reward zone was activated. Alternation training continued until rats reached asymptote performance (rat 1 = 27 days and rat 2 = 9 days; rat 3 = 20 days). The data reported here for rats #1 and #2 comes from the next, *random lights,* task, in which sequences of up to 900 elements were drawn randomly with replacement from the 32 light/reward zones, with each LED remaining active until the rat reached it. For rats #1 and #2, 25 recording sessions and 23 recording sessions were obtained, respectively. A different random sequence was used for each behavioral session to ensure that learning effects did not contributing to the activity patterns. The data reported here for rat #3 comes from the last, spatial sequence, task that followed training on the random lights task. For the spatial sequence task, rats were trained to navigate through an 8-item repeating element sequence of reward zones (**Fig. 1**). This task consisted of 3 traversals through the sequence with cue lights illuminated immediately (cued) and 3 traversals with a 5s delay (delay-cued), thus demonstrating sequence memory (Bower et al., 2005). For delay-cued trials, trained rats completed the trial before the appearance of light cues ≥ 90% of the time on average.

**Figure 1.**
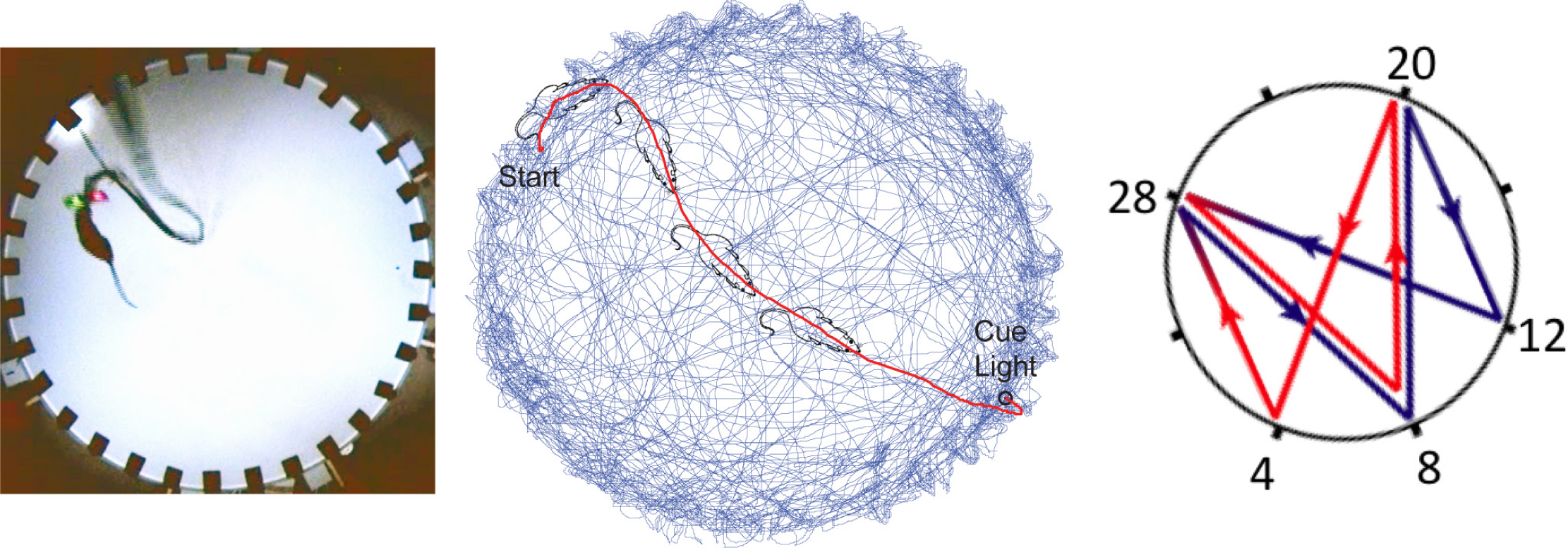
Apparatus, reference frames, and learned motion sequence. *Left.* Apparatus. Rats #1 and #2 were trained to run a random spatial sequence to 32 light locations. This task requires the rat to learn to execute a stereotyped motion sequence, and covers the full range of headings and body-centered cue light directions at a wide range of spatial locations. *Middle.* Single segment from the random lights task **(red)** overlaid on 99% transparent subset (1/5th) of the trials from that session **(blue)**. Note, that in the later training phases the route to the goal becomes a series of stereotyped motion sequences. *Right. Schematic of the spatial sequence task.* The rat starts at zone 12 and continued to zone 28 8 20 4 28 8 20 & 12. Rat #3 learned to execute this spatial sequence from memory (without cueing).

Rats #1-2 were housed in a vivarium with lights on between 07:30 and 19:30 and were tested during the light cycle. Rat #3 was housed in a reverse light/ dark cycle vivarium and tested during the dark cycle. For rat #3, prominent distal cues were arranged around the perimeter of a large room (~4.5 m x 6 m). For the remaining rats, prominent cues (strips of white curtain) were displayed on 1-2 walls of a square curtain that hung ~1m beyond the edge of the apparatus. For all rats and sessions, dim ambient light illuminated the maze from above. All experiments were carried out in accordance with protocols approved by the University of Lethbridge Animal Welfare Committee and conformed to NIH Guidelines on the Care and Use of Laboratory Animals.

### Recording Procedures

A custom electrode interface board attached to the recording array with independently drivable tetrodes connected 3 unity-gain headstages (HS-27, Neuralynx) to the recording system (Neuralynx). Tetrodes were referenced to an electrode in the corpus callosum and advanced as needed, up to 60μm/day, while monitoring the audio and visual signal of the unit activity, but adjustments were carried out only after a given day’s recording to allow overnight stabilization. Once a large number of units in the parietal cortex were obtained, alternation training commenced. Thresholded (adjusted prior to each session) spike waveforms were filtered 0.6 to 6 kHz and digitized at 32 kHz. A continuous trace was simultaneously collected from one of the tetrode wires (filtered 0.1 to 9 kHz, and digitized at 2034.75 Hz and referenced to an electrode in corpus callosum) for processing as a local field potential (LFP) and timestamps were collected for up to 18 tetrodes. Rat position and head direction were tracked using colored domes of reflective tape, which were created by covering [$] Styrofoam balls with reflective tape (Fig. 1), and on-line position information was used to trigger MFB stimulation rewards. Position and head direction data were collected at 60Hz as interleaved video (rats 1 & 3) or 30Hz (rat 2) and co-registered with spikes, LFPs, and stimuli.

Spike data were automatically overclustered using KlustaKwik then manually adjusted using a modified version of MClust (A.D. Redish). All spike waveforms with a shape and across tetrode cluster signature suggesting that they were likely multi-unit activity (MUA) and not noise, were manually selected and merged into a single MUA cluster. Thus, MUA clusters included both well-isolated single units and poorly isolated single units. MUA clusters were not treated as unique unless a) the tetrode was moved > 100 μm from the previous session and b) the distributions of spike clusters were clearly unrelated between the two sessions (http://klustakwik.sourceforge.net; Harris et al., 2000).

LFP analyses were performed using custom-written Matlab code (Mathworks, Nattick, MA) or Freely Moving Animal (FMA) Toolbox (http://fma-toolbox.sourceforge.net/). The LFP signal was collected at 2034.75Hz and subsequently resampled to 2000 Hz for further analysis, using the Matlab resample function. The amplitude of the high frequency (HF) LFP signal was obtained by digitally filtering the raw signal in the 300-900 Hz range using a 4-th order zero-phase Chebyshev filter and calculating the absolute value of the Hilbert transformed filtered signal.

### Parietal Cortex Data

Data recorded from a subset of the sessions (47 sessions) included MUA clusters that were assessed because the tetrode had moved at least 100 μm from the previous session. Data from these 47 sessions (rat #1 = 14 sessions, rat #2 = 18 sessions, rat #3 = 15 sessions) included 176 potentially unique MUA clusters (number of putative unique MUA clusters for rat #1 = 46, rat #2 = 55, and rat #3 = 75). Each recording session (except for 12 sessions of the 47 total sessions presented here) consisted of three 50 min rest sessions intermixed with two 50 min behavioral sessions on the apparatus. The remaining 12 sessions consisted of one behavioral session between two rest sessions, followed by a 4h break during which the animal was returned to the vivarium before returning for a second round of a 50 min behavioral session between two more 50 min rest sessions. For these 12 sessions only 1 of the rest-task-rest sessions was analyzed for a given day.

### Self-Motion Analyses

Position and head direction data were utilized to map the self-motion reference frame for each MUA cluster. For these analyses, position data were interpolated and smoothed by convolution with a Hamming window that was 1s long. In addition, head direction data gaps <1s were transformed using directional cosines for interpolation using the interp1 function in Matlab, then transformed back to polar coordinates (Gumiaux et al., 2003). Next, head angular velocity, linear velocity and MUA activity rate were calculated for each video frame using a 100ms sliding window. Finally, the occupancy and number of MUA spikes for each 3 cm/s by 20°/s bin were calculated and converted to firing rate for each bin with >0.5 s of occupancy. For illustrative purposes (not cell classification analyses), the self-motion activity rate maps were smoothed by convolving with a Gaussian function for the 2 x 2 bins surrounding each bin (Chen et al., 1994a). The self-motion colormaps represented a range of MUA (or HF) activity rates from 0 **(blue)** to the maximum **(maroon)**. No adjustments were made to the standard, evenly spaced colormap. MUA clusters were classified as having a preferred-self motion state if the common points with sufficient occupancy (i.e., >0.5 s) from the self-motion maps for the first and second daily session (or split ½ were significantly positively correlated (p<0.01). This was generally the most conservative criterion for self-motion cells of the three criteria reported by Whitlock et al. (Whitlock et al., 2012). Specifically, for each MUA cluster, to determine if cells had “significant” self-motion properties, the map from the first daily behavioral session was shuffled, a correlation coefficient was computed between the first session (shuffled map) and the second session (unshuffled map), and this process was repeated 500 times. Then, the second behavioral session map was shuffled, the correlation coefficient was computed between the second session (shuffled map) and the first behavioral session (unshuffled map), and this process was repeated 500 times (total 1000 shuffles/cell). The entire shuffled data set for each cell was used to calculate a critical r-value for the 99th percentile. For the day recording sessions with two behavioral sessions, stability was assessed by comparing across the two sessions and for the five recording sessions where a single behavioral epoch was available, split ½ measures of stability were used (i.e., a single session was split in ½ based on time and the first ½ was compared to the second ½).

### Memory Replay Analyses

First, still periods were extracted from the rest sessions as described previously (Euston et al., 2007; Johnson et al., 2010). The raw position data from each video frame was smoothed by convolution of both x and y position data with a normalized Gaussian function with standard deviation of 120 video frames. After smoothing, the instantaneous velocity was found by taking the difference in position between successive video frames. An epoch during which the velocity dropped below 0.78 pixels/s for more than 2 minutes was judged to be a period of motionlessness. All analyses of rest sessions were limited to these motionless periods. Slow wave sleep (SWS) and rapid eye movement (REM) sleep were distinguished using automatic K-means clustering of the theta/delta power ratio extracted from the CA1 pyramidal layer LFP recorded during the ‘stillness’ periods (Girardeau et al., 2009). Sharp-wave ripples (SWR) were detected from the CA1 LFP digitally filtered in the 75-300 Hz range. Events with peaks > 5 standard deviation (SD) above the mean and duration less than 100 ms were considered SWRs. The SWR duration included the contiguous periods surrounding the peak and exceeding 2 SD above the mean. The SWR detection accuracy was visually validated on the subset of each analyzed dataset.

#### Template Matching Memory Replay Analyses

We performed *template matching* analysis, previously used to show the simultaneous reactivation of isolated single neuron ensembles in the medial prefrontal cortex (Euston et al., 2007; Louie and Wilson, 2001a). The criteria for the inclusion of a dataset in the template matching analysis was at least 12 min of SWS and 600 SWRs during both pre- and post-task-sleep, and at least 50 trials during the task. Template matrices (number of tetrodes x number of time bins; **Fig.2A**) were generated from the trial-averaged multiunit activity (MUA template) or Hilbert-transformed high frequency amplitude (HF template), extracted from a 2 s/trial windows that preceded the arrival at the reward site, and binned to 100 ms bins. Only trials longer than the template duration (2 s) were included. The time window was chosen based on evidence that reactivation of the hippocampal activity patterns is more prominent for the task phase immediately preceding the reward (Diba and Buzsaki, 2007; Foster and Wilson, 2006; McNamara et al., 2014; Singer and Frank, 2009). The time bin choice was based on previous reports of template matching using isolated single unit neuron activity (Euston et al., 2007; Johnson et al., 2010), where a 100 ms bin was deemed optimal for capturing task-related neuronal dynamics. In order to eliminate tetrodes with sparse and/or poorly approach-motion-modulated activity, only the tetrodes with average MUA > 1 Hz and binned spike train coefficient of variation > 0.25 during reward approach period were included in the template, which resulted in elimination of 0-17% of tetrodes.

**Figure 2.**
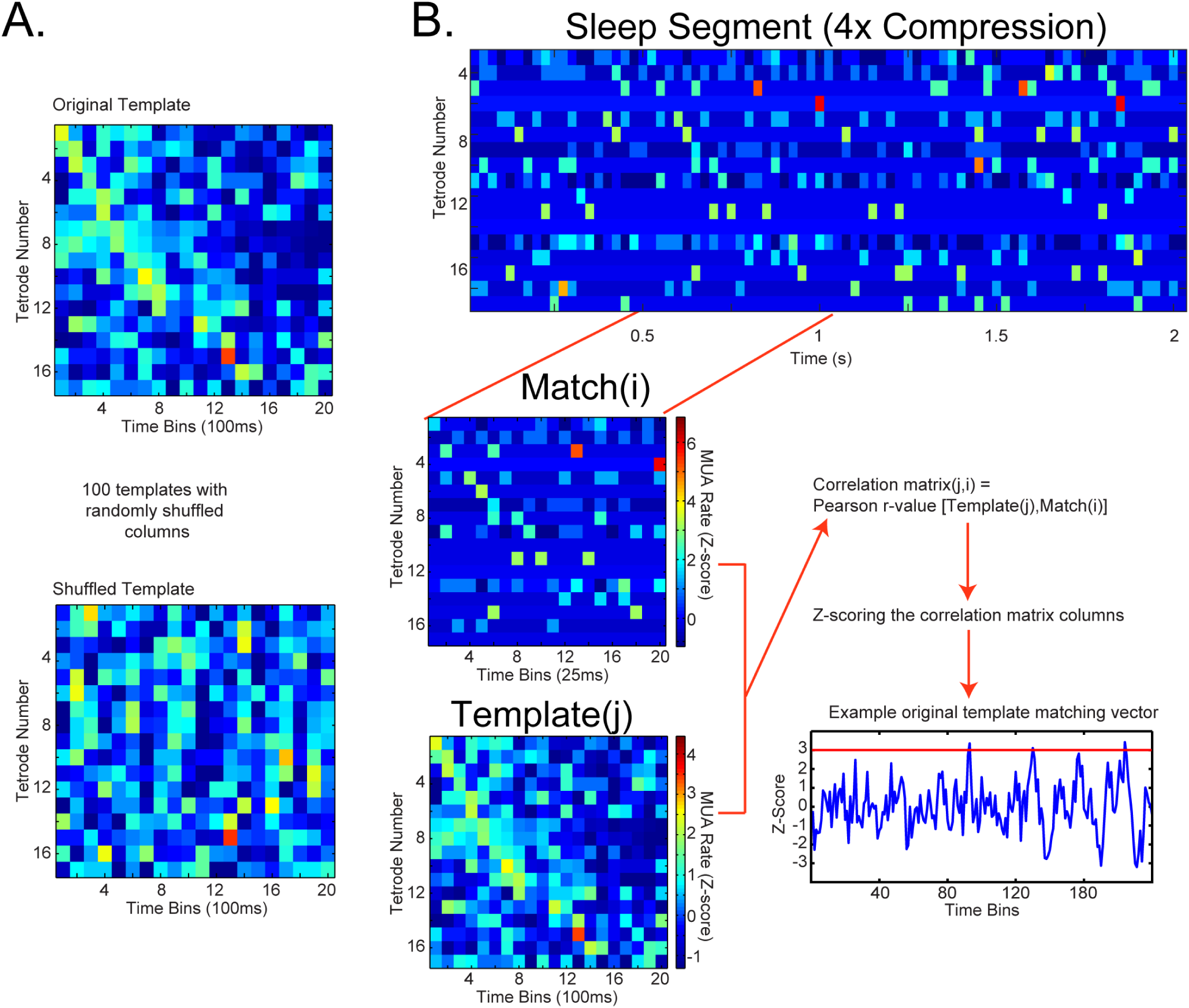
Template matching method. **A.** Shuffling procedure. Above. Example original template. Multiunit activity (MUA) or high-frequency (HF) amplitude on each tetrode (row) is averaged over all the trials, binned at 100 ms and Z-scored. Below. Template shuffling procedure. Position of each column (instantaneous MUA or HF amplitude over all the tetrodes) was randomly permuted, in order to produce 100 shuffled templates while preserving the overall MUA or HF levels, as well as the instantaneous correlation between tetrodes. **B.** Template matching procedure. *Above*. MUA data segment from post-task sleep. Note, firing during sleep is sparse, so samples are more variable. Activity on each tetrode (row) was binned at a bin size of 100 ms / compression factor and Z-scored. *Below Left*. Example match window from the sleep epoch (above) and template (below). Pearson correlation coefficient was calculated between the template and equally sized slow-wave sleep (SWS) segments of the sleep session, which were produced by sliding the template-size window over the sleep epoch with a 1-bin step size. This creates the correlation matrix (number of templates x number of time bins during sleep epoch). Each column of the correlation matrix is Z-scored, the Z-score value reflecting the degree of similarity of a given template to sleep activity at given sleep time window, relative to the distribution across the original and all the shuffled templates. *Bottom right*: Example original template matching trace. Z-score values above 3 are considered matches.

Only the templates containing 6 or more tetrodes were retained in the analysis. To allow comparison between the MUA and HF template matching, all the HF templates were constructed from the same set of tetrodes that passed the MUA-based screening. To eliminate the influence of the MUA or HF amplitude variability between tetrodes, the binned signal was Z-scored for each tetrode separately.

MUA and HF amplitude from the sleep periods was processed in the same way, except that the bin size was adjusted according to the compression factor (bin size = 100 ms / compression factor; e.g. for the 4x compression, the sleep bin size was 25 ms), to capture the compressed nature of neural reactivation during sleep. An evenly spaced range of compression factors (4x, 6x, 8x, 10x), as well as the ‘no-compression’ (nc) control were used. Only slow wave sleep periods were included in the analysis.

To test the matching of a given template and the pattern of activity during sleep *(matching significance),* each template was shuffled repeatedly to generate 100 shuffled templates. The shuffling procedure consisted of randomly permuting the position of each column in the template (population vector), preserving the overall activity levels and instantaneous correlations between the MUA or HF signals on different tetrodes, but scrambling the sequential patterns. A Pearson correlation coefficient was calculated between each template and the series of candidate matches, generated by sliding the template-size window over the sleep epoch (**Fig. 2B**). This resulted in a matrix of Pearson correlation coefficients r, where the element r^i,^j corresponded to the correlation coefficient between the i-th template and j-th candidate match. The correlation matrices were Z-scored across individual time bins (columns), and the resulting Z-score values reflected the template similarity to the corresponding sleep segment at given time step, relative to the distribution that included the original and 100 shuffled templates. Z-score values above 3 were considered *matches*. The threshold for *matching significance* was that the number of matches from the original template had to exceed the number of matches from all of the 100 shuffled templates for a given sleep epoch (i.e., p < 0.001).

For comparison of template matching between the pre- and post-task-sleep, match percentage was obtained by dividing the number of matches by the number of SWS time bins for each sleep epoch. For the ripple-triggered average analysis, the original template Z-score traces +/- 1 s around each SWR peak time were averaged, obtaining the ripple-triggered averaged Z-score for a given sleep epoch.

#### Statistical Analyses

Two-way repeated measures ANOVA (both variables are repeated measures) was used to assess the main effect of sleep session (pre-versus post-task sleep), the main effect of compression factor (nc, 4x, 6x, 8x, and 10x), and interactions (sleep session x compression factor) on template matching measures. Significant interactions were followed up by planned comparisons (F-tests) comparing pre-task versus post-task-sleep for each compression factor. Except where noted otherwise, p < 0.05 was considered statistically significant. For example, one exception was for self-motion map correlations, for these analyses we used the historical critical value of 0.01 (Whitlock et al., 2012; Wilber et al., 2014).

#### Histology

After the final recording session, rats were deeply anesthetized with Euthasol and transcardially perfused with 0.1M phosphate buffered saline followed by 4% paraformaldehyde. The whole head was post-fixed in 4% paraformaldehyde with electrodes in place for 24h, then brains were extracted and cryoprotected in 30% sucrose. Frozen sections were cut (40μm) using a sliding microtome or custom block-face imaging system (Leica vibratome), mounted on chrome alum-subbed slides, stained with cresyl-echt violet, and imaged using a NanoZoomer Imaging system (Hamamatsu).

## RESULTS

Results from MUA and HF-LFP analyses were largely similar. Therefore, we report the MUA results here and provide the corresponding HF-LFP data and some comparisons in Supplemental materials.

### Modular Tuning to Motion State

We looked for evidence of population level encoding in egocentric coordinates in the rat parietal cortex. To do this, we assessed MUA recorded on single tetrodes as a function of either head direction, egocentric cue light location, or linear, and angular velocity in rats that had been trained to either run 1) the cued random spatial sequence (rats 1 & 2) or 2) a complex repeating element spatial sequence from memory (rat 3). In contrast to our previous single unit analysis we did not find evidence for MUA tuning to head direction (Chen et al., 1994a; Wilber et al., 2014), and we did not find evidence for MUA tuning to egocentric cue light location (not shown). It is possible the lack of head direction tuning for MUA but presence with single cells means that the head direction tuned cortical columns are present but are very narrow and thus not detected with MUA. Next, we determined if parietal cortex MUA had a preferred self-motion state (e.g., right turn) by plotting firing rate as a function of the rat’s current linear and angular velocity. MUA was classified as having a preferred self-motion state if the self-motion rate maps for two behavioral sessions were significantly positively correlated (i.e., self-motion rate maps; Chen et al., 1994a; Whitlock et al., 2012; Wilber et al., 2014). We found that the MUA recorded on a single tetrode was frequently significantly tuned to a specific self-motion state (i.e., specific combination of linear and angular velocity). In fact, nearly every tetrode was significantly tuned for at least one recording depth (41 of 44 tetrodes; 93%) of the tetrodes that were positioned in parietal cortex for at least one session.

Interestingly, the tuning appeared to be invariant across depth. To quantify this observation, we collected all of the pairs of sessions where the tetrode was at least 100 μm from the comparison location and the self-motion maps for both sessions were significant (p<0.01). For each of these pairs we tested the two maps for significant similarity versus a random shuffle distribution using the same method as for within-day behavioral sessions. Interestingly, self-motion tuning appeared to be consistent across cortical laminae because tuning was consistent on a single tetrode when recordings were compared even for very large spacing between recording depths (**Fig. 3B**). In fact, a large majority of comparisons were significant (**Fig. 3C;** 73%), indicating significant correlation in modular self-motion tuning across a range of cortical depths for a particular tetrode. In addition, as a control we performed depth correlations across tetrodes (instead of within tetrode) in the same manner (i.e., we took each pair of sessions that was separated by at least 100 μm) and found significantly more correlations with an r-value < 0 (χ2_(1)_=5.09, p<0.05) and significantly fewer significant cross-tetrode map correlations (χ2_(1)_=5.68, p<0.05). As an additional control we compared the separation of significant and non-significant depth spacing for comparison pairs. The distribution of depths for significant versus non-significant depth correlations was highly overlapping: significant Mean ± SEM = 280 μm ± 25 and non-significant Mean ± SD = 241 μm ± 31. There was no significant difference in the difference in depths between the two samples for significant depth correlations (t_(72)_=0.85, p=0.40), suggesting that closer spacing was not responsible for the significant depth correlations.

**Figure 3.**
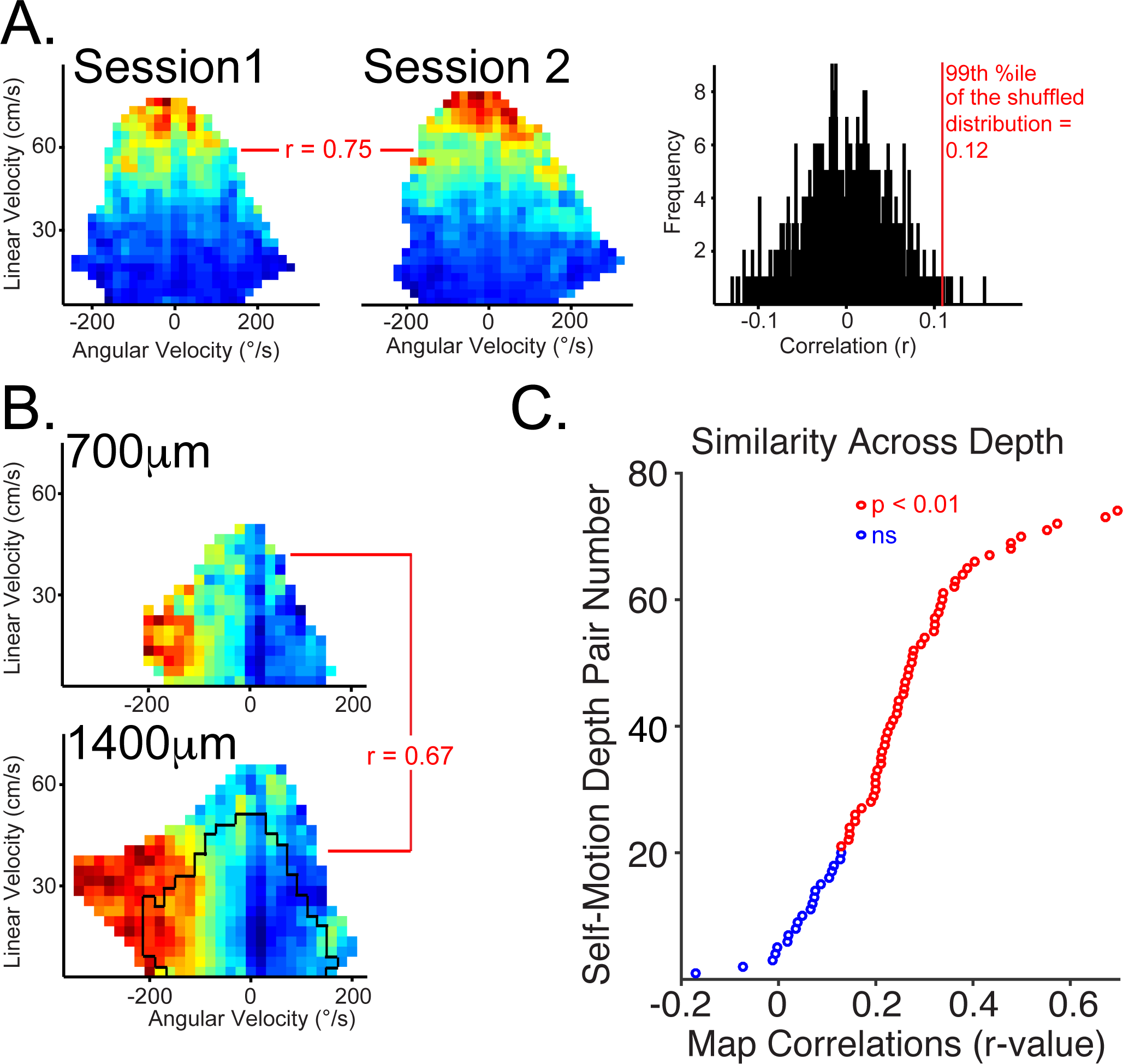
Self-Motion tuning in parietal cortex is invariant across cortical laminae. See also Figure S1. **A.** *Left Two Plots.* Multi-unit activity recorded for a single day’s recording session and from a single tetrode were classified as having a preferred-self motion state if the self-motion maps for two behavioral sessions (from the whole day recording session) were significantly positively correlated. Self-motion maps from two behavioral sessions and corresponding correlation value are shown for one multi-unit module. Occupancy data and number of spikes are binned according to linear velocity (vertical axis) and head angular velocity (horizontal axis; positive head angular velocity corresponds to a right turn), then converted to activity rate (number of multi-unit spikes per second). *Right.* The shuffled distribution and critical r-value corresponding to the 99th percentile is shown. The map from the first behavioral session was shuffled 500 times and a correlation coefficient was computed between the shuffled and unshuffled maps. Then, this process was repeated to shuffle the map for the second session 500 times (total 1000 shuffles) in order to calculate a critical r-value for the 99th percentile (p<0.01). See Materials and Methods for additional details. **B.** Same as in A; however, data came from two separate recording sessions obtained when the tetrode was at two different depths (700μm, *above* and 1400μm, *below*). Black outline on lower motion rate map illustrates that for cross-depth comparisons behavior can vary considerably, and this analysis is limited to common data points. **C.** The sorted correlation value for each MUA cluster for each pair of depths where the tetrode was moved at least 100 μm and the session data for each depth met the significance criteria described in A. Pairs of depths with significantly correlated motion maps (as described in B, p<0.01) were colored **red** indicating significant correlation in modular self-motion tuning across a range of cortical depths for a particular tetrode. Data comes from all tetrodes that met this criteria from all 3 rats.

Finally, if we substituted the amplitude of the HF-LFP envelope (300-900Hz) for MUA firing rate, similar tuning maps were observed and were often significantly correlated with the corresponding MUA map (**Fig. S1**), consistent with the theory that the HF-LFP signal reflects neuronal spiking. Further, when we took each data set with significant within-session MUA motion map stability and compared the correlation values for single recording session (behavioral session 1 vs behavioral session 2 maps) obtained using HF-LFP signal to the correlation values obtained when generating the same maps with MUA, we found that these correlation values are significantly correlated (r=0.29, p<0.01). It should be noted that although correspondence can be quite good for these two measures and often subjectively appears very impressive (as shown in **Fig. S1**), during behavior at least, the HF signal tends to produce poorer tuning on average. We quantified this slight difference by taking all of the self-motion maps that were ‘significant’ at two different thresholds (p<0.05 and p<0.01) and comparing the number of instances where a self-motion map is significant at p<0.05 (but not p<0.01) versus significant at the higher p<0.01 level for HF vs MUA. Significantly more MUA data sets were significant at the higher (p<0.01) level (χ2_(1)_=5.03, p<0.05).

### Patchy, modular topographical organization of motion correlates

In contrast to the high degree of laminar consistency of behavioral correlates, the topographic organization was patchy and modular: sometimes, a tuning pattern was consistent across several near-by sites, but there were also abrupt transitions between some neighboring sites. Illustrations of this organization are provided in Figure 4.

**Figure 4.**
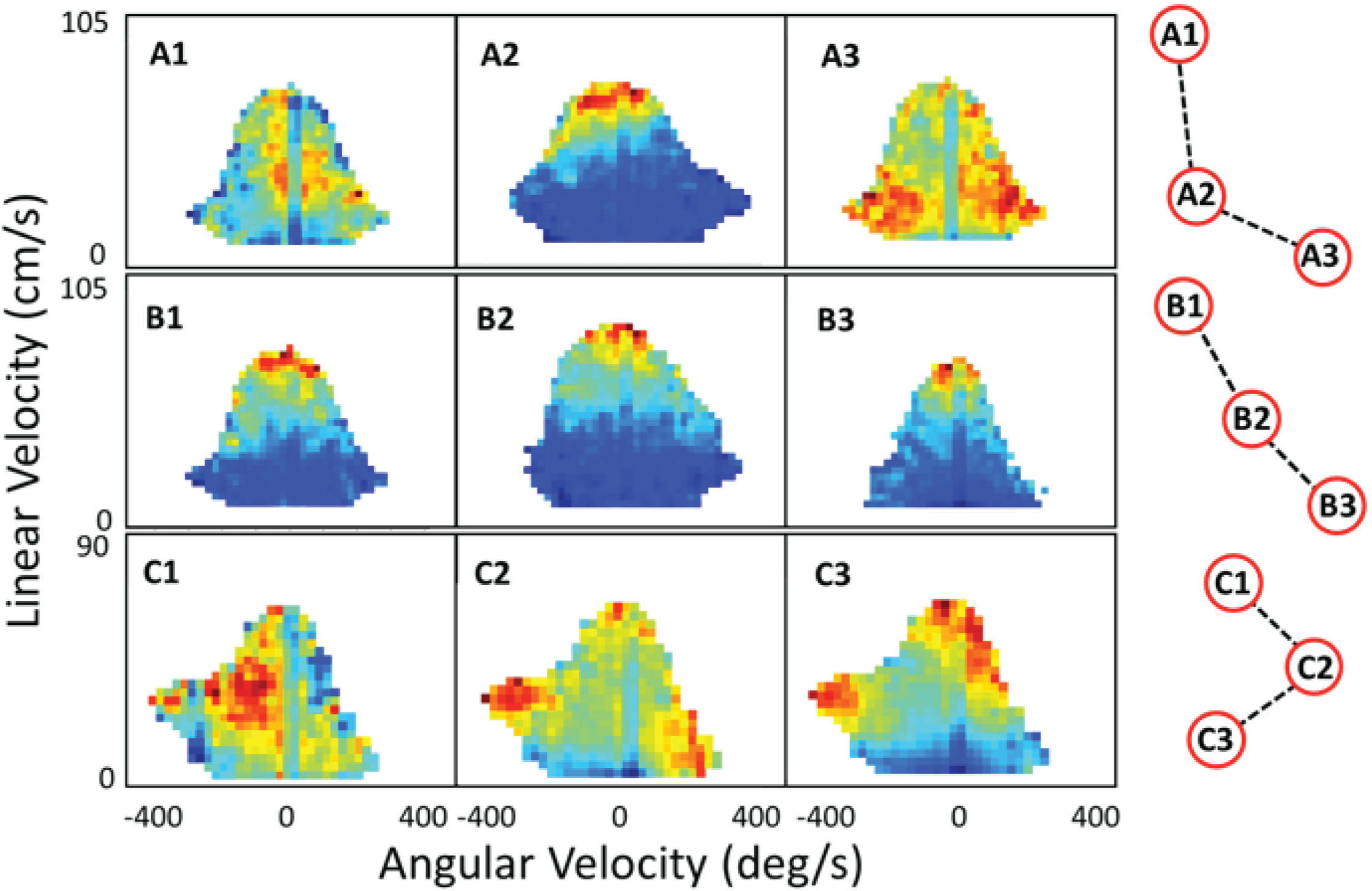
Illustration of patchy modularity of MUA behavioral correlates in parietal cortex. Rows A and B represent behavioral correlate maps for three nearby electrodes in left and right hemispheres respectively from rat 1. Row 3 is from the left hemisphere of rat 2. The relative positions of the electrodes are shown on the right, where the minimum distance represented is about 300 μm. In Row A, the correlates change abruptly from position to position, whereas in the Row B, the correlates are quite similar from position to position. Row C illustrates a combination of these effects. As shown in Figure 3, the correlates were highly consistent in the laminar dimension for each location. For illustration, the session with the most significant behavior map was selected for each electrode. Thus, sessions were different for the examples shown in A and B (and as a result the map shape varied), but was the same for the examples shown in C.

### Modular Interactions During Sleep

The random lights task required the animals to execute a series of actions in a relatively stereotyped manner (i.e., a motion sequence), something the rats learn to do over the course of training (see **Fig. 1** for an example of a path that has become stereotyped). Thus, we set out to explore the possibility that this learned behavior may be reflected at the neural level in the form of memory reactivation involving self-motion modules during sleep periods.

### Template Matching

#### Parietal modular activation patterns during slow wave sleep resemble patterns observed during the task

All the template matching analysis was limited to SWS periods. First, we compared modular parietal activation patterns from the task to post-task-sleep. We analyzed results from 15 data sets (5 data sets from each rat), selected based on distribution across training, numbers of trials (>50), presence of sufficient quantity of slow-wave sleep (> 12 min / sleep epoch), number of sharp wave ripples (>600 / sleep epoch) and number of tetrodes in parietal cortex that met criteria for inclusion in the template (>5). A majority of these data, 12 data sets, had 3 sleep epochs (pre-, mid- and post-task-sleep interleaved with two task sessions) while the remaining 3 data sets had two sleep epochs (pre- & post-task-sleep with 1 task session). Consistent with previous reports, we found that memory reactivation was more consistently stronger for the post-task-sleep session in data sets with two task and three sleep epochs (Euston et al., 2007; Johnson et al., 2010). Therefore, we restricted our analysis to these data sets (12 sets of pre-task-sleep, task, and post-task-sleep from 3 rats). Due to the time-compressed nature of re-activation reported previously for single cell memory reactivation studies (Euston et al., 2007; Peyrache et al., 2015; Wilson and McNaughton, 1994), we performed template matching for several evenly spaced compression factors: no-compression, 4x, 6x, 8x, and 10x. The sleep sessions during which the original approach template had significantly larger number of matches (ps < 0.001) relative to shuffled distribution was considered significantly matching. The 10x compression produced the lowest (12/24, 50%), while the compression factor with largest proportion of significantly matching sleep epochs was 4x (19/24, 79%). For the HF amplitude template the lowest proportion of significantly matching sleep epochs was found with no-compression (20/24, 83%), while the largest proportion was found with 4x and 10x compression (23/24, 96%). Overall, the proportion of significantly matching sleep sessions was higher for HF amplitude behavioral templates.

In order to test the hypothesis that modular template matching during sleep is a part of memory consolidation process and does not simply reflect the reverberation of certain activity patterns during sleep, we compared the match percentages for ‘no compression’ and range of equally spaced compression factors (4-10x), between the pre-task-sleep and post-task-sleep. For the MUA templates, post-task sleep had significantly higher template matching percentage (percentage of time bins with a match that exceeded Z-score value of 3) than pre-task sleep (**Fig. 5A**; main effect of sleep session, F_(1, 11)_ = 11. 34, p < 0.01) and this effect varied across compression factors (sleep session x compression factor interaction; F_(4, 44)_ = 11.18, p < 0.0001). In addition, there was a main effect of compression factor (F_(4, 44)_ = 6.97, p < 0.001). Planned comparisons were conducted for pre-versus post-task-sleep for each compression factor. Post-task-sleep had significantly more template matches for 6x, 8x, and 10x (Fs_(1, 22)_ > 4.73, p < 0.05) but not 4x (Fs_(1, 22)_ = 3.98, p = 0.058) compression factors, while pre- and post-task-sleep match percentage did not differ for ‘no-compression’ (F_(1, 22)_ = 0.75, p = 0.40).

**Figure 5.**
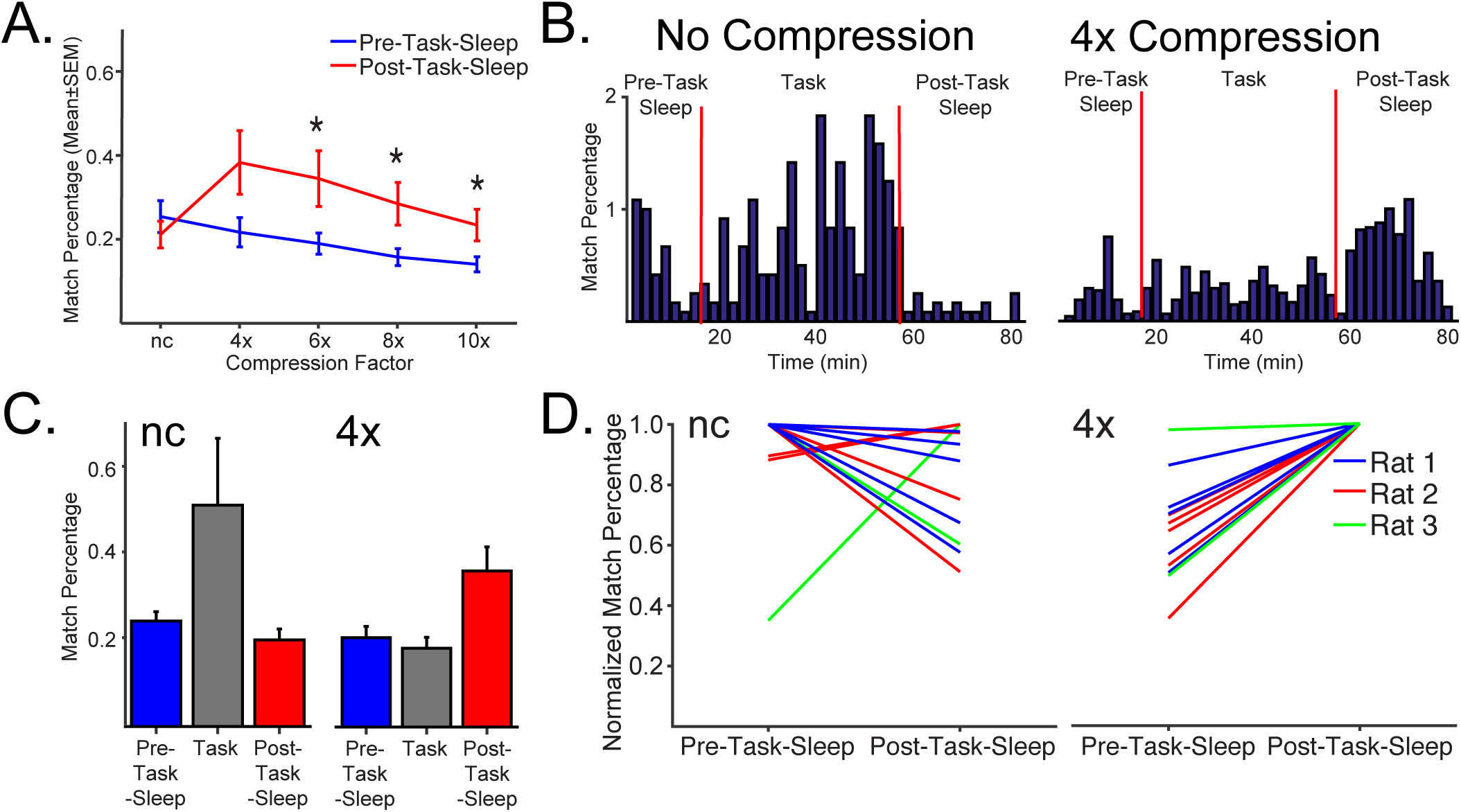
Modular sequence reactivation occurs in parietal cortex and is compressed. See also Figure S2. **A.** Mean (±SEM) match percentage (number of matches/number of time bins) across the compression factors for multi-unit activity (MUA,) templates for slow-wave sleep periods. Template matching increases between pre- (blue) and post-task-sleep (red) for compressed data, but not for ‘no-compression’. For MUA and HF amplitude templates, reactivation measures peak at 4x compression. **B.** Example of MUA showing the match percentage (2 min bins) for ‘no-compression’ (no-compression, left) and 4x compression (right), over the pre-task-sleep, task and post-task-sleep. For ‘no-compression’, there is a high match percentage during the task, and very low in post-task-sleep. For the 4x compression (right), match percentage is much higher in post-task-sleep, relative to both pre-task-sleep and task. Only the SWS periods from each sleep epoch are shown. **C.** Mean of the pre-task-sleep, task and post-task-sleep as was calculated for each session and averaged across time bins that are shown in B. Then a mean for all sessions was calculated for each rat. Finally, the mean (±SEM) of the rat mean data is shows that across rats reactivation is stronger in post-task rest when a 4x compression factor is applied, but not for ‘no compression’. To avoid the possible contribution of awake reactivation to template matching during task, only the contiguous movement periods (>5 cm/sec) longer than 2 sec were used in quantifying the template matching during task. **D.** Normalized match percentage (number of matches / number of time bins divided by the peak value for that session) across the ‘no-compression’ (left) and 4x (right) compression factors for multi-unit activity (MUA) templates for slow-wave sleep periods for each session for each of 3 rats. Reactivation consistently increases between pre- and post-task-sleep for compressed data, but not for ‘no compression’. For MUA and HF amplitude templates, reactivation measures peak at 4x compression. Datapoints are normalized to pre-task-sleep values. * p<0.05.

Similar to MUA-based reactivation measures, HF amplitude post-task sleep had significantly higher template matching than pre-task-sleep (**Fig. S2**; main effect of sleep session, F_(1, 11)_ = 21.51, p < 0.001) and this effect varied across compression factors (sleep session x compression factor interaction; F_(4, 44)_ = 9.20, p < 0.001). In addition, there was a main effect of compression factor (F_(4, 44)_ = 21.51, p < 0.0001). Planned comparisons for pre- versus post-task-sleep for each compression factor showed that for ‘no-compression’ pre-task-sleep did not significantly differ from post-task-sleep (F_(1, 22)_ = 1.47, p = 0.24), while post-task-sleep had significantly more matches than pre-task-sleep for 4x – 10x compression factors (Fs_(1, 22)_ > 5.87, ps > 0.05).

As illustrated for the MUA template example in **Figure 5B**, for ‘no-compression’, there is a high match density during the *task*, and low match density in *post-task-sleep*. A dramatically different pattern is shown for 4x compression, where the match percentage was much higher in *post-task-sleep*, relative to both *pre-task-sleep* and *task*. Next, the mean of the pre-task and post-task-sleep was calculated for each session and averaged across the time bins shown in **Figure 5B**. Then a mean for all sleep and task sessions was calculated for each rat, for all the data like the example shown in **Figure 5B**. To avoid the possible contribution of awake reactivation to template matching during task, only the contiguous movement periods (>5 cm/s) longer than 2 s were used in quantifying the template matching during task. This showed that reactivation strength for pre- vs post-task-sleep varied dramatically across compression factor (‘no compression’ & 4x compression; **Fig. 5C**) as a function of task phase. First, post-task-sleep was greater than pre-task-sleep but only for 4x compressed data. Second, the matches where much higher during the task phase for ‘no compression’ data than for 4x compression data, reflecting the fact that the templates do match well to the behavior session that was used to generate them. Stronger reactivation during post-task rest was not just observed for the session means (**Fig. 5A**), individual time bins for individual data sets (**Fig. 5B**) and the rat means (**Fig. 5C**), but also when the pre-versus post task rest normalized (to post-task rest) was plotted for ‘no-compression’ versus 4x compression data for each session for each rat (**Fig. 5D**). Nearly identical results were obtained for HF amplitude (**Fig. S2**). Overall, MUA template – based reactivation measures increased in post-sleep.

#### Parietal cortex reactivation is increased during hippocampal sharp wave ripples (SWRs)

Finally, we explored the role of hippocampal-cortical interactions in modular memory reactivation by quantifying the strength of parietal cortex template matches as a function of SWR events in the hippocampus. We found that the strength of the SWR-triggered template match was greater during post-task sleep for MUA based templates (**Fig. 6A**), and also for HF amplitude based templates (**Fig. S3**). For both MUA and HF amplitude templates, the peri-SWR reactivation strength (maximum averaged ripple-triggered Z-value within +/- 500 ms peri-SWR time-window) was significantly greater during post-task-sleep for (**Fig. 6B**; main effect of rest session; Fs_(1, 11)_ > 16.69, ps < 0.01), and the significant effect of rest session varied across compression factors (Fs_(4, 44)_ > 3.01, ps < 0.05). The only difference between measures was that there was a significant main effect of compression factor for HF amplitude templates (**Fig. S3B**; F_(4, 44)_ = 18.87, p < 0.001), but not MUA templates (F_(4, 44)_ = 3.01, p = 0.25). To follow-up on the significant interaction for MUA and HF amplitude data, planned comparisons for pre-versus post-task rest were conducted for each compression factor. The ripple-triggered Z-score amplitude was significantly larger for all compressed data (4x-10x; Fs_(1, 22)_ > 4.77, ps < 0.05), but not for ‘no-compression’ (Fs_(1, 22)_<1.97, ps>0.17). Thus, for both MUA and HF signal templates the amplitude of ripple-triggered Z-value was significantly larger for post-task-sleep particularly for intermediate compression factors.

**Figure 6.**
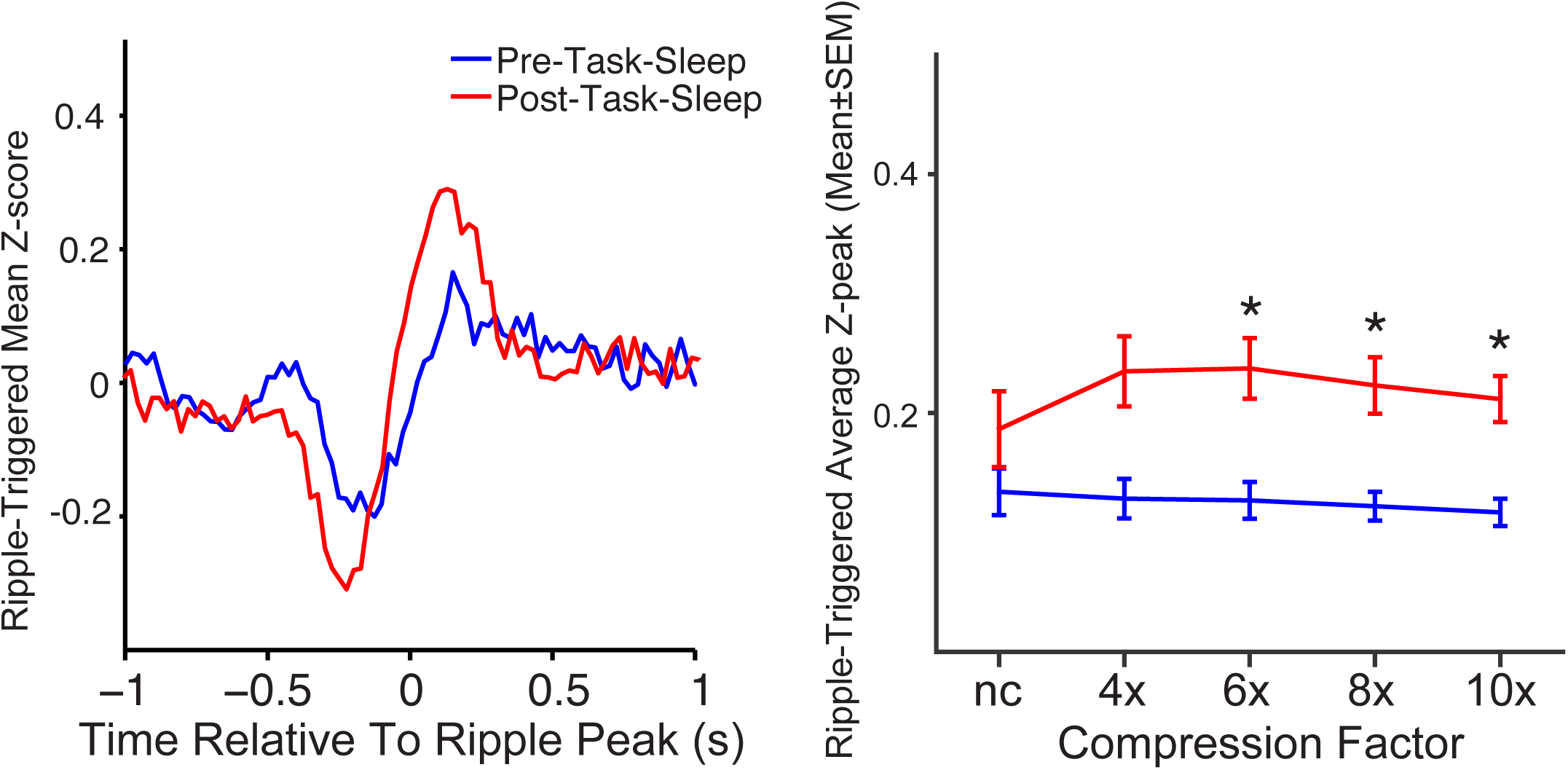
Modular sequence reactivation in parietal cortex is enhanced around sharp-wave ripples (SWRs) in the hippocampus. See also Figure S3. ***Left.*** Example of sharp-wave ripple (SWR) triggered average Z-scores for *pre-task-sleep* **(blue)** and *post-task-sleep* **(red)** with 4x compression. SWR triggered peak amplitudes for multi-unit activity (MUA) templates, decaying to baseline within 300-400ms after ripple peak. ***Right***. Ripple-triggered average peak amplitudes across the compression factors for MUA templates. Peak amplitudes varied significantly across compression (F_(4, 44)_ = 3.01, p < 0.05). Only slow-wave sleep periods were included in the analysis. * p<0.05.

#### Sleep parameters

To assess the possible differences in sleep architecture between pre-task-sleep and post-task-sleep we compared the durations of different sleep states (SWS, REM), as well as their ratios across these sleep epochs. There was not a significant effect of sleep epoch on the duration of REM sleep or REM/SWS ratio (ts_(11)_<0.98, ps>0.35; **Table 2**). SWS duration was significantly greater in post-task-sleep (ts_(11)_>2.23, p<0.05); however, match percentage reflects the number of matches/time unit so more slow-wave sleep will not impact this measure.

**Table 1.** Training and Testing Conditions. Recording sessions refers to daily sessions that were typically split into two 50 min behavioral sessions separated by a 50 min rest session.

| Rat # | Task Type | Light/Dark Cycle Phase During Testing | Number of Recording Sessions |
| --- | --- | --- | --- |
| 1 | Up to 900-item fully random | Light | 25 |
| 2 | Up to 900-item fully random | Light | 23 |
| 3 | Spatial Sequence Task | Dark | 16 |

**Table 2.** Sleep architecture metrics for pre-versus post-task-sleep. For the REM duration and REM/SWS ratio, there were not significant differences in pre-versus post-task-sleep (ts_(11)_<0.98, ps>0.35). However, SWS duration was significantly greater in post-task-sleep (SWS duration: t_(11)_=2.23, p<0.05).

|  | Pre-Task-Sleep | Post-Task2-Sleep |
| --- | --- | --- |
| **SWS duration (s)** | **1850 +/- 647** | **2484 +/- 349** |
| **REM duration (s)** | **141 +/-37** | **200 +/- 226** |
| **REM/SWS ratio** | **0.09 +/- 0.08** | **0.09 +/- 0.11** |

## DISCUSSION

We have shown modular coding of movement in parietal cortex. Further, the coding is consistent across a range of cortical depths, strengthening the idea that this meso-scale coding is organized in a potentially modular fashion. In addition, we found that there are interactions between modules during post-task-sleep as measured by templates reflecting the task segments leading up to the reward zone. These modular memory reactivation events suggest that inter-modular interactions are involved in normal learning and memory. Finally, we demonstrated that the high frequency LFP amplitude signal from a given tetrode produces a result that is highly similar to the multi-unit activity for both modular motion tuning and memory reactivation.

Single unit recording studies and behavioral studies have pointed towards an egocentric reference frame in parietal cortex. However, this is the first demonstration of mesoscale functional encoding from a local population of single cells organized around a specific motion preference. For example, coordinate transformation networks may exist embedded within a functional module for executing the appropriate action sequence (Byrne et al., 2007; McNaughton et al., 1995; Wilber et al., 2014; Xing and Andersen, 2000). In fact, the motion modules could possibly even serve as a common output reference frame to a brain network for decision related computations involving parietal cortex (Goard et al., 2016; Hanks et al., 2015; Harvey et al., 2012; Licata et al., 2016). In other words this modular movement based encoding may be a fundamental coding framework in parietal cortex. This common tuning, likely reflecting high intrinsic connectivity among the single units (i.e., a module; Buxhoeveden and Casanova, 2002) suggestive of a cell assembly. Such a functional module may be precisely the effector unit theorized to be critical for sufficient size to trigger responses in target areas (Breitenberg and Schulz., 1991; Bush and Sejnowski, 1994; Gabbott et al., 1987) either at the column (Mountcastle, 1997; Szentagothai, 1975) or possibly even macrocolumn level (Mountcastle, 2003).

It should be noted that we did not examine all known encoding modes in parietal cortex. One encoding framework that seems likely to exist at the modular level is route-centered encoding, described in detail at the single unit level (Nitz, 2009; Nitz, 2006, 2012). Unfortunately, the task we employed here is not optimal for detecting route-centered modulation. In contrast to our previous single unit analysis that suggested consistent head direction tuning across depth, we did not find evidence for MUA tuning to head direction (Chen et al., 1994a; Wilber et al., 2014). It is possible the lack of head direction tuning for MUA but presence with single cells means that the head direction tuned cortical columns are present but are very narrow (e.g., microcolumn) and thus not detected with MUA. In addition to head direction encoding other types of world-centered (allocentric) encoding have been reported in parietal cortex that could exist at the modular level but were not assessed here (Nakamura, 1999). Similarly, outside of parietal cortex, there are numerous examples of local cell populations that share functional similarities, including primary visual areas, and the barrel cortex (Petersen, 2007; Roudi et al., 2015; Sincich and Horton, 2005). Local populations with functional similarity also exist in higher cortical areas, for example, grid cells in the medial entorhinal cortex (Moser et al., 2014; Roudi et al., 2015) or object feature columns in inferior temporal cortex (IT; Tsunoda et al., 2001). Presumably the current approach can be applied to further understanding of many of these brain systems.

We demonstrated modular reactivation in parietal cortex at the ensemble levels, using template matching. This method was previously applied in memory reactivation studies of ensembles of isolated single neurons (Euston et al., 2007; Louie and Wilson, 2001b; Pavlides and Winson, 1989; Skaggs and McNaughton, 1996; Tatsuno et al., 2006; Wilson and McNaughton, 1994). Although the resemblance between the behavioral and sleep activity patterns could be due to simple re-occurrence of the similar patterns across the sleep-wake cycle, multiple lines of evidence support the notion of modular memory reactivation in the present study. First, the behavioral template matching was more prevalent during post-task, relative to pre-task sleep, an effect present after normalizing for the SWS duration in each sleep epoch (**Fig. 5**). This suggests that the difference in probability of certain brain activity patterns between pre- and post-task-sleep is affected by intervening experience, one of the critical requirements for the detection of memory-related processes. Second, consistent with numerous reports of memory reactivation occurring at the temporally compressed timescale (Euston et al., 2007; Nadasdy et al., 1999; Peyrache et al., 2009; Wilson and McNaughton, 1994), we observed increased matching during post-task-sleep for the range of compression factors (4-10x) while no significant change was observed for non-compressed patterns. Temporal compression during memory consolidation is postulated to create the optimal conditions for spike-timing dependent plasticity (Lansink and Pennartz, 2014; Markram et al., 1997). Finally, temporal coupling of modular reactivation in parietal cortex with SWRs suggests that, similar to reactivation in medial pre-frontal cortex (mPFC; Euston et al., 2007; Peyrache et al., 2009), this process is part of hippocampo-cortical communication during sleep, one of the hallmarks of memory consolidation (Maingret et al., 2016).

We found a certain degree of significant matching with pre-task-sleep activity, consistent with previous reports (Dragoi and Tonegawa, 2011; Peyrache et al., 2009). This is consistent with the idea that the information incoming during experience modifies the pre-existing patterns (Luczak et al., 2009), but does not completely perturb them. The reactivated patterns resemble the templates constructed based on the average activity during approach to reward locations randomly assigned on each trial, and therefore likely encode the behavioral sequence reinforced by subsequent reward. Reactivation of behavioral pattern averaged over many different spatial trajectories, containing a range of head direction, velocity and other tuning properties of parietal cortex, supports the abstraction role of memory consolidation (Lewis and Durrant, 2011; Tse et al., 2007), as the regular aspects of episodic memory are extracted into the semantic domain.

Finally, we showed that the HF-LFP signal has nearly identical reactivation dynamics compared to dynamics observed with MUA activity. Although the exact contribution of LFP generation mechanisms to different frequency ranges are a matter of debate (Buzsáki et al., 2012), correlation between the HF amplitude and MUA activity is highly correlated with the level of spiking activity recorded at the individual electrode, which suggests that the instances of low correlation between MUA and HF amplitude are due to spike subsampling on the individual electrode. Another factor that could decrease the MUA-HF amplitude correlation is that the spiking contribution to HF amplitude depends on the individual neuron type and recent spiking history, as during bursting activity, where the amplitude of subsequent spikes tend to decrease. Finally, detection of spikes that comprise MUA clusters were thresholded and thus these clusters represent spiking activity within a more limited spatial range than HF amplitude which is not thresholded and thus could potentially detect more distant spiking activity.

Consistent with numerous reports describing mostly cortical-hippocampal interactions (notably mostly the pre-frontal cortex), we found enhanced reactivation of behavioral templates simultaneously with SWRs in the CA1 field of the hippocampus (Jung et al., 1998; Peyrache et al., 2009; Peyrache et al., 2015; Siapas and Wilson, 1998; Singer and Frank, 2009; Wierzynski et al., 2009). Though a recent report found that equal numbers of prefrontal cortex single cells were excited or inhibited during hippocampal SWRs; while other single cells were not modulated by the SWR at all (Peyrache et al., 2009). This suggests that the modular re-activation we observed may represent a subset of the single cells in that cortical region.

In summary, we have shown modular self-motion tuning in rat parietal cortex. These motion modules also participate in memory re-activation as measured by templates constructed from the pre-reward task phase. Our findings suggest a potential fundamental level of organization in the rat in parietal cortex and open up a new avenue for bridging the gap between macro-level measures of brain activation in humans and unit recording studies in non-human animals.

## Author Contributions

AAW and BLM designed the experiment; AAW conducted the experiment; AAW and IS analyzed the data; AAW, IS and BML wrote the paper.

## Acknowledgements

We thank Mikko Oijala, Dr. David Euston, Dr. Michael Eckert, and Dr. Masami Tatsuno for technical assistance with analyses and Scott Killianski and Christin Montz for thoughtful comments as we constructed analyses. Alberta Innovates Health Solutions Polaris Award to BLM. AG049090, MH4682316, and an Alberta Innovates Health Solutions Fellowship to AAW.

